# Precision data-driven modeling of cortical dynamics reveals idiosyncratic mechanisms underlying canonical oscillations

**DOI:** 10.1101/2023.11.14.567088

**Authors:** Matthew F. Singh, Todd S. Braver, Michael W. Cole, ShiNung Ching

## Abstract

Task-free brain activity affords unique insight into the functional structure of brain network dynamics and is a strong marker of individual differences. In this work, we present an algorithmic optimization framework that makes it possible to directly invert and parameterize brain-wide dynamical-systems models involving hundreds of interacting brain areas, from single-subject time-series recordings. This technique provides a powerful neurocomputational tool for interrogating mechanisms underlying individual brain dynamics (“precision brain models”) and making quantitative predictions. We extensively validate the models’ performance in forecasting future brain activity and predicting individual variability in key M/EEG markers. Lastly, we demonstrate the power of our technique in resolving individual differences in the generation of alpha and beta-frequency oscillations. We characterize subjects based upon model attractor topology and a dynamical-systems mechanism by which these topologies generate individual variation in the expression of alpha vs. beta rhythms. We trace these phenomena back to global variation in excitation-inhibition balance, highlighting the explanatory power of our framework in generating mechanistic insights.

## 1. Introduction

A key goal of human neuroscience is to decipher how individual differences in brain signaling and dynamics relate to individual differences in cognition and behavior. Developing mechanistic models of individual human brains is one part of this endeavor. While considerable efforts have been directed at identifying individual differences at macroscopic spatial scales, via fMRI, individual-differences in fast dynamical interactions at the scale of M/EEG, while well-documented, are less understood. Such dynamics reveal neural computation at a timescale commensurate with sub-second cognitive operations and are strongly nonstationary. There is a long and rich history of analysis of fast neural electrophysiology, yet the mechanisms and functional salience of many commonly observed phenomena remain debated. A notable example of such ambiguity concerns canonical EEG oscillations, including the posterior dominant alpha (8-12 Hz) rhythm. Generative mechanisms of alpha oscillations have been studied for decades, but are not yet resolved, with competing accounts of either a thalamic [1, 2] or cortical origin [3]. At a phenomenological level, alpha tends to vary across individuals in terms of its peak frequency and power [4, 5, 6], and furthermore is associated with various cognitive endpoints [7, 8, 9]. As a result, it is a frequent candidate as a biomarker [10, 11], including to inform brain stimulation strategies [12]. Such implementations, which are largely empirical in nature, underscore the need for reliable, biologically-plausible dynamical systems models with sufficient expressiveness so as to reveal individual mechanistic differences. Despite a century of research on the alpha rhythm, there have been few results and no consensus regarding why healthy individuals differ in alpha expression [13]. To preview, we develop a modeling framework which sheds new light on this debate, by providing mechanistic insights and generating testable predictions regarding the nature of alpha, as well as other oscillatory individual difference phenomena.

### 1.1. Challenges in Data-Driven Individualized Modeling

Dynamical systems models are premised upon describing how components of a system interact to shape its future. The classical example of such models in neuroscience is, of course, the Hodgkin-Huxley [14] model of neuronal spiking. The power of these models is that they are simultaneously descriptive and mechanistic. That is, they not only describe features of the observed phenomena but also provide an underlying generative process that produces those phenomena. This property means that dynamical systems models can potentially predict how the system will respond to novel perturbations. This ability is based on the accuracy of the underlying model, which in turn depends upon how the model is constructed.

Data-driven approaches to dynamical systems modeling attempt to use measured brain activity to parameterize (i.e., ‘fit’) a model. At the whole-brain scale, there have been significant efforts directed towards this problem in the context of functional neuroimaging. These previous approaches to individualized brain modeling include methods that directly estimate parameters from fMRI recordings, such as Dynamic Causal Modeling [15] or, alternatively, methods in which the primary model parameters (e.g., connectivity) are adopted from structural imaging (e.g. [16, 17, 18]). In the former, the prime difficulty has been the development of methods to estimate large, non-linear models from noisy, indirect timeseries. In recent work, we developed a new algorithm termed Mesoscale Individualized NeuroDynamic modeling (MINDy, [19, 20]) to estimate individualized brain models from fMRI timeseries. The general method consisted of a novel optimization framework to simultaneously estimate brain network parameters and, in an extended algorithm [20], local hemodynamic responses. However, the temporal resolution of fMRI greatly limits its ability to inform models of the fast, transient interactions thought to dominate neural computation. In the current work we aim to reveal these mechanisms at the individual level and are therefore concerned with high-temporal resolution modalities. As our approach consists of whole-cortex modeling, we emphasize the use of MEG or EEG (M/EEG) as functional data-sources, as opposed to lower-coverage invasive techniques (e.g., ECoG, LFP, and SEEG), although the proposed method is general. In either case, the fast timescale context produces a new set of challenges for individualized modeling that evade current optimization methods, including the original MINDy framework [19].

There are theoretical, biophysical, and computational barriers to this endeavor. At the theoretical level, fast electrophysiological activity, such as oscillations, are hypothesized to arise from the interplay of excitatory and inhibitory neurons [21, 22], meaning that any biologically interpretable model of brain oscillations must consider the interactions between specific neuron types including long-distance projections onto either cell-type. In addition, asymmetric patterns of connectivity (feed-forward vs. feed-back) are also believed critical to generating lower-frequency oscillations [23]. However, neither of these two features is accessible using structural (diffusion) imaging. Functional data is similarly limited in directly assessing these differences using conventional analysis of M/EEG signals. Both modalities are thought to be driven by cortical pyramidal cell activity as interneuron geometry is not conducive to dipole generation [24]. From an optimization (model-fitting) standpoint, these limitations mean that neither the model states (excitatory and inhibitory neural activity) nor model parameters are directly accessible, posing a challenging dual-estimation problem. To address these difficulties, we present a new framework to directly estimate detailed neural-mass style models (Fig. 1A) from fast functional data (M/EEG). We term this framework Mesocopic Individualized NeuroDynamics with Dual Estimation (Dual MINDy). To be clear, by ‘directly estimate’ we mean optimizing every component of the neural model to predict observed activity (timeseries measurements), within subject. The net result of our framework will encompass: (i) the estimation of latent activity in neural populations across the cortex, (ii) separate brain-wide ‘connectomes’ for excitatory and inhibitory targets (Fig. 1A), and (iii) direct model-estimation from single-subject recordings.

**Figure 1.**
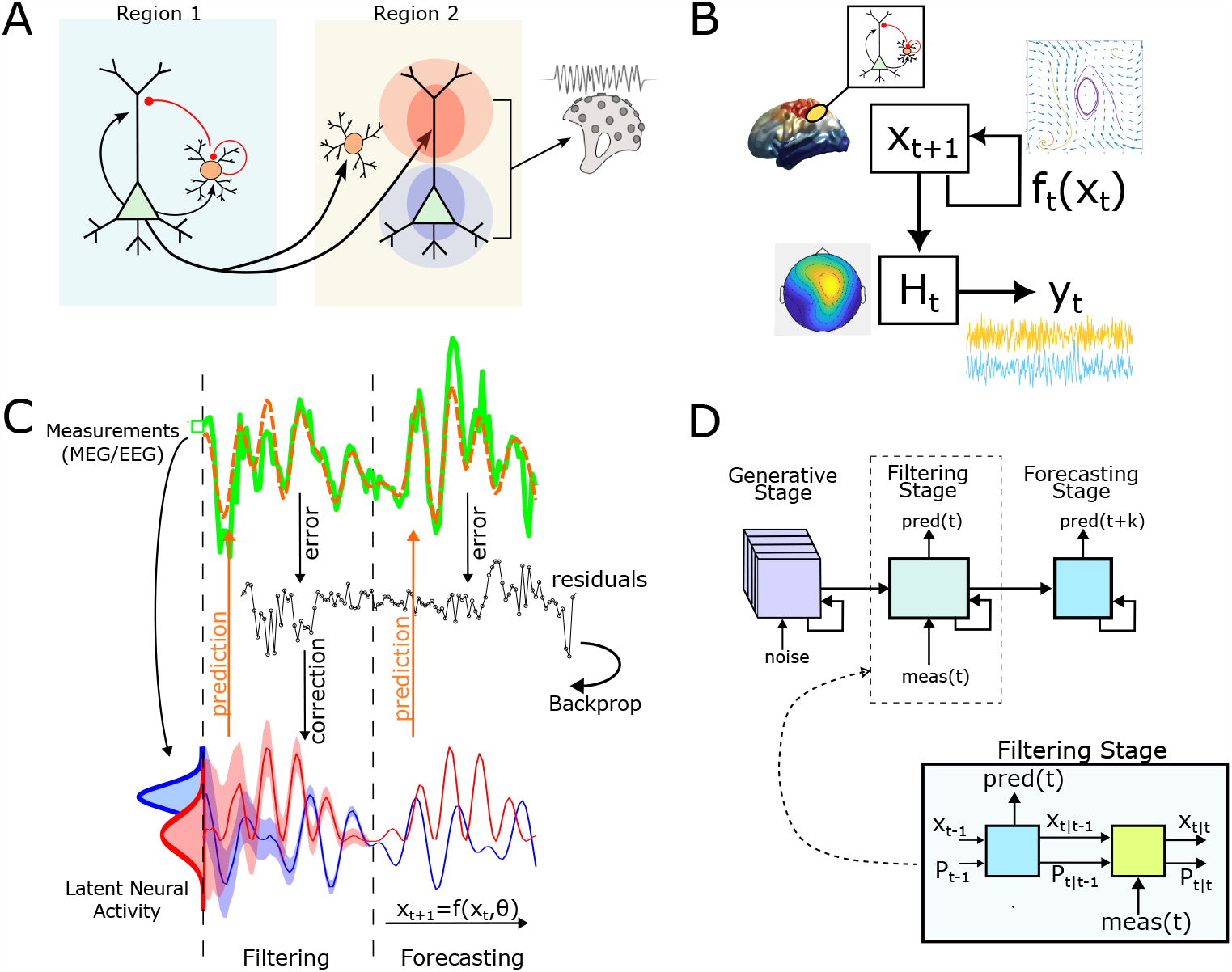
Schematic description of the proposed framework. A) Expanded neural-mass type model with long-distance connections onto both excitatory and inhibitory targets originating from distal pyramidal cells. Local EI circuits are fully connected. Arrows/dots indicate (directed) termination site. Dipoles are modeled as proportional to excitatory (pyramidal) depolarization at the some. B) Relationship between components of the combined state-space and measurement models. Neural activity (*x*(*t*)) evolves according to the dynamics *f* (*x*). Measurements (*y*_*t*_) are produced by multiplying neural activity with the measurement matrix *H*_*t*_. C) The gBPKF algorithm consists of three stages: generating baseline distributions, a Kalman-Filtering stage to estimate latent states (red and blue), and a Forecasting stage which predicts future brain activity measurements (green). Distributions (bottom left) indicate the posteriors at *t*_0_. Note that the uncertainty (shading) decreases over time as the (nonlinear) Kalman filter corrects state-estimates. D) The algorithm instantiated as a nonlinear recurrent network. The generative (noise-driven) and forecasting (deterministic) layers evolve as conventional recurrent networks, whereas the filtering stage uses the Kalman filter to evolve both activity/states and uncertainty/covariance. Steady-state distributions from the generative stage are used to estimate the initial state/uncertainty for *t*_0_.

We will proceed to introduce the Dual MINDy framework and validate it on the Human Connectome Project (HCP; [25]) dataset. Furthermore, we will highlight the mechanistic explanatory power of the method in the context of cortical oscillations, by studying individual variability in generative processes underlying M/EEG oscillations. Such oscillations are among the most frequent features extracted from M/EEG, yet their underpinning and cognitive significance remains an open question, in part due to their variable expression across individuals. Here, we show that this variation may reflect low-dimensional dynamics that we link with a greater ratio of excitation-to-inhibition. We suggest that individual variation in alpha-band frequencies are reflected in protracted, global rhythms, whereas variation in the beta-band is linked to transient dynamics.

## 2. Results

### 2.1. Dual MINDy enables scalable, data-driven cortical modeling

We seek a large-scale cortical model of the mesoscopic, mean-field type:

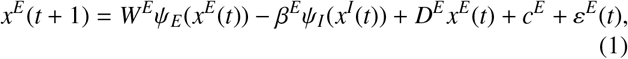

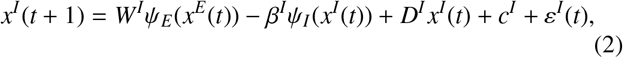

This model is, in essence, a network of interconnected Wilson-Cowan type neural masses [26], each modeling excitatory-inhibitory interaction at the scale of cortical macrocolumns. Here, *x*^*E*^ and *x*^*I*^ describe the activation (average depolarization) of excitatory and inhibitory subpopulations, respectively, and the nonlinear function *ψ* is a 2-parameter logistic function with gains (*s*^*E*^,*s*^*I*^) and bias terms *v*^*E*^, *v*^*I*^. Full technical details regarding the model are found in the Supplemental Information (SI).

Our goal, simply stated, is to optimize (i.e., fit) all free parameters (see Table 1) of this model on the basis of observed brain activity. To do so requires the formulation of a measurement model that transforms *x*^*E*^, *x*^*I*^ into sensor ‘outputs’ *y*_*t*_. Here, we assume *y*_*t*_ is acquired from a noisy transformation *H* of underlying neural activity (Fig. 1B):

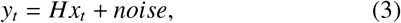

where *x* ≡ [*x*^*E*^, *x*^*I*^]. The formulation (3) leads to a crux issue for data-driven model parameterization in this context. Specifically, for M/EEG signals, the measurement matrix *H* is not invertible, even with accurate source-localization, because *x*^*I*^ does not contribute directly to the M/EEG signal. Hence *H* takes the form: [*H*_*Exc*_ 0_*k*×*n*_] (for *k* measurement channels and *n* populations). The transformation of excitatory activity *H*_*Exc*_ could be direct (using the lead-field matrix for *H*_*Exc*_) or, if data is already source-localized, *H*_*Exc*_ = *I*_*k*=*n*_. However, in either case, the unknown latent activity of all populations *x*_*t*_ is not directly recoverable from *y*_*t*_. This leads to the need for *dual-estimation*, encompassing the combination of two-problems: estimating states *x*_*t*_ and identifying parameters. Such problems are quite challenging in any circumstance. The application to brain-network modeling further challenges existing dual-estimation approaches, which become computationally-intractable due to the large number of unknown parameters [27], primarily in terms of network connections *W*^*E*^, *W*^*I*^.

**Table 1:** Summary of free parameters.

| Parameter | Interpretation |
| --- | --- |
| $W^E, W^I$ | Exc.-Exc. and Exc.-Inh. connectivity |
| $1-D^E, 1-D^I$ | Exc. and Inh. decay rates |
| $\beta^E, \beta^I$ | Local Inh.-Exc. and Inh.-Inh. connections |
| $c^E, c^I$ | Tonic drive to exc. and inh. populations |
| $\varepsilon^E, \varepsilon^I$ | Extrinsic (unmodeled) activity |

Our framework derives from the observation that both halves of this problem are individually well-studied and tractable, but cannot be applied in isolation (estimating *x*_*t*_ requires knowledge of parameters, and vice-versa). Instead we remove the problem of estimating latent brain-activity by replacing *x* with a pseudo-optimal estimate given all parameters and previous measurements: 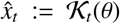 (where *θ* denotes the parameters), which produces a conventional parameter-estimation problem (solve for *θ*, Fig. 1C). In other words, rather than treating states and parameters as unknown variables, we first define the best estimate of state, given parameters, and then solve for parameters which optimize this function. In practice, we use nonlinear-variants of the Kalman filter [28, 29] for the state estimate and attempt to minimize the prediction-error with respect to future measurements *y*_*t*+*k*_. In short, we solve for parameters that generate the most-accurate Kalman filter. To retrieve these parameters, we treat the Kalman-filter recursions like a recurrent network (Fig. 1D) and analytically backpropagate error gradients through the entire algorithm (see SI Sec. 8.4, Fig. 1C) which we combine with gradient/Hessian optimization. This technique, which we term as a generalized Backpropagated Kalman Filter algorithm (gBPKF, [30]; also see related earlier work by [31]), has been validated and shown to be scalable for general classes of circuit models, but not specifically validated for the meanfield form (1)-(2) in the presence of biophysical constraints. Again, full details pertiaining to the gBPKF are found in the SI.

Our first set of analyses focused upon determining whether we could reconstruct all connectivity parameters in *W*^*E*^, *W*^*I*^, containing both recurrent and long-distance connections, given the observed timeseries with all other parameters known. We generated ground-truth, simulated networks with either 20 or 40 regions each containing an excitatory and inhibitory population (40 or 80 total populations). Only excitatory populations generate long distance connections. The symmetric connection graph of admissible connections had a sparsity of 25% nonzero with the same graph used to define admissible EE and EI connections. Connection strengths were not symmetric in either case. The ratio of simulated channels to regions (75%) was based on the empirical rank of leadfield matrices (typically 65-80) relative the 100 parcels we later use.

Results demonstrate high performance in recovering ground-truth connectivity parameters in biologically-plausible simulations (Fig. 2A). We observed high-performance in recovering the true connectivity in both the 40 population (*EE* : *r* = .983 ± .005, relative-MSE:.021 ± .006; *EI* : *r* = .919 ± .048, 60 sims) and 80 population conditions (*EE* : *r* = .970 ± .004, relative-MSE:.029 ± .003, *EI* : *r* = .884 ± .014, 30 sims). The performance for excitatory-excitatory connections was particularly strong (Fig. 2A). This advantage is expected, as the excitatory populations directly contribute to the simulated M/EEG signal whereas the effects of EI signaling are only indirectly observed through their delayed propagation along local IE coupling. However, despite this challenge, performance remained high. We conclude that our gBPKF algorithm is well-suited to estimate neural model parameters for all connectivity types (EE, EI, etc.).

**Figure 2.**
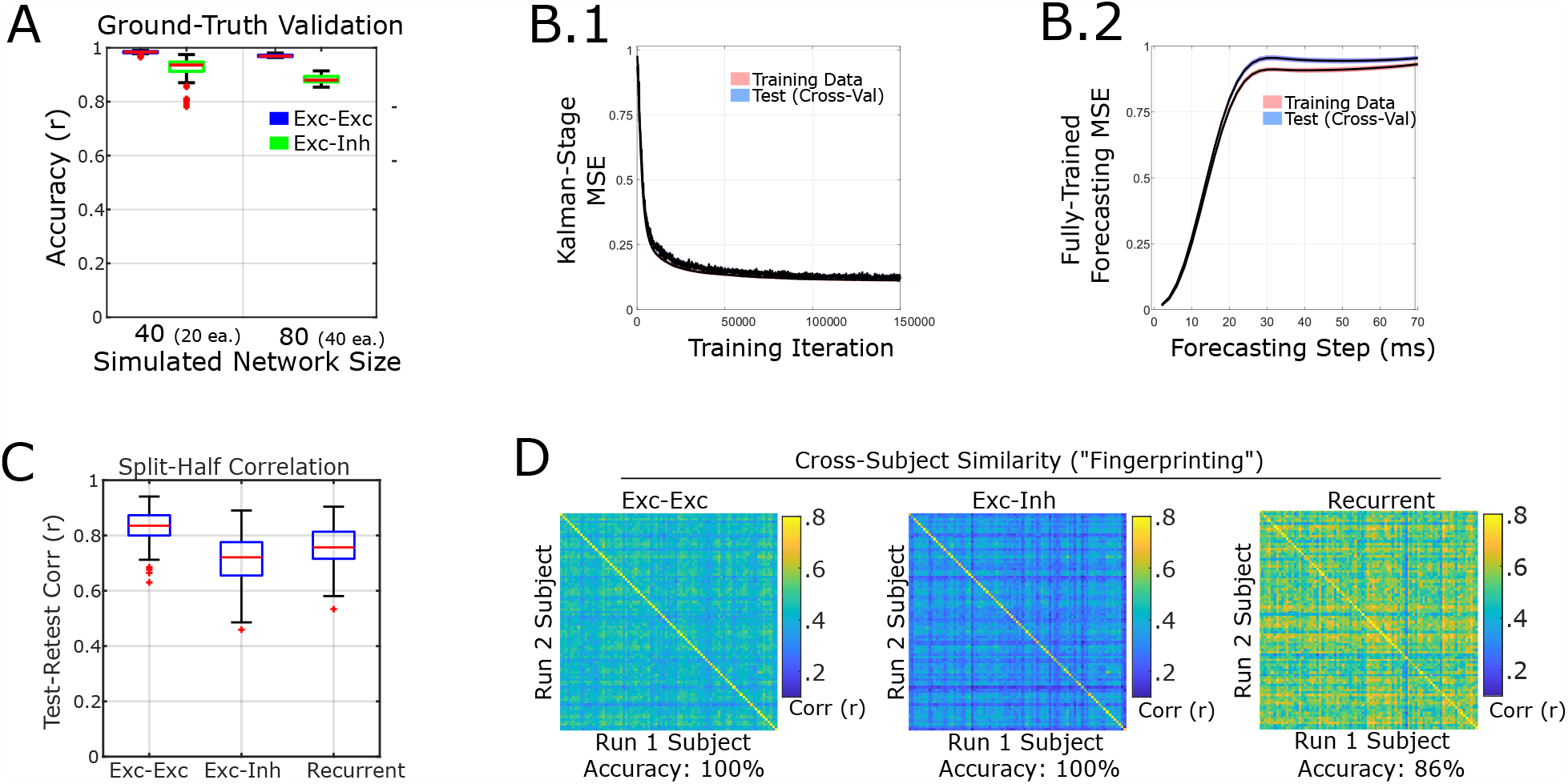
Validation of the current framework. A) Our method accurately recovers the excitatory-excitatory and excitatory-inhibitory connection strengths in realistic ground-truth simulations containing either 40 or 80 total populations (20 or 40 regions x 2 populations per region). B) Forecasting error converges and indicates similar performance on training and left-out data, meaning that models are stable and generalizable to new data within-subject. B.1) Change in error during the Kalman Filtering phase across training iterations during model estimation. B.2) Performance in forecasting future brain activity for various lags after 150k training iterations. Lines indicate the mean loss across subjects/runs and shading indicates standard error of measurements. Shading indicates ±1 standarderror of measurement (n=174) for both (B) panels. C) Connectivity parameters are reliable across models trained on different data for the same subject. D) Model parameters are individual-specific, forming a unique “fingerprint” [32] that matches parameters fit to different data from the same subject. Accuracy indicates the percent of successful identifications (i.e., how often two models from the same subject are most similar, as opposed to another subject’s model).

### 2.2. Models provide reliable estimates of individual brain dynamics

Next, we fit models to the HCP MEG data, which contains three five-minute runs per subject. We divided this data into chronological halves (seven minutes each) which we refer to as a “scan”, We first analyzed the reliability of model parameters. For univariate parameters, we measured reliability in terms of the Intra-Class Correlation (ICC) which assesses reliability of individual differences. For multivariate parameters we present both the conventional test-retest correlations for overall similarity of the parameter and, for the reliability of individual differences, the Image Intra-Class Correlation (I2C2, [34]) which is a multivariate extension of ICC.

As in our ground truth simulations, the primary parameters of interest for reliability are the connectivity parameters. Combined together, the connectivity matrix is highly reliable and individualized. Excitatory-excitatory long-distance connections had exceptional test-retest correlations (*r* = .83 ± .07, *I*2*C*2 = .72; Fig. 2C). Long-distance excitatory-inhibitory connections also had good test-retest correlations (*r* = .71 ± .1) but were more modest for individual differences (*I*2*C*2 = .46). We also found good reliability for the spatial gradient of recurrent connections (*r* = .75 ± .10). Due to co-dependency (see SI Sec 8.7), the test-retest correlation, but not the I2C2 is the same for all recurrent connection types. Individual differences in recurrent connections were higher for inhibitory targets than excitatory: I2C2: *EE* = .58, *IE* = .56, *EI* = .85, *II* = .86. Fingerprinting accuracy was 100% for both EE and EI connectomes (Fig. 2D) and 84% for the spatial-gradient of recurrent connections.

Individual differences in decay rates had good reliability for excitatory populations (*ICC* = .70) and moderate for inhibitory populations (*ICC* = .64). The nonlinear connection gain (*S* ^*E*^) also had good reliability (*ICC* = .72). Estimated noise standard deviations 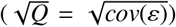 often saturated (lower-bound 0.1, upper-bound 0.3), and were, therefore, poor markers for individual differences 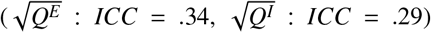. As expected (SI Sec. 8.7), reliability was low for the non-linear threshold (*v*^*E*^ : *I*2*C*2 = .36) and baseline drive terms (*c*^*E*^ : *I*2*C*2 = .35, *c*^*I*^ : *I*2*C*2 = .12). While the inclusion of these parameters is important for reproducing the correct dynamics, they contribute via their values relative each-other (after a transformation) and, depending upon the forward model, may not be individually unique (see SI Sec. 8.7 for discussion). Group-average values for connectivity parameters and the spatial gradient of recurrent connections are displayed in Fig. 3A-C.

**Figure 3.**
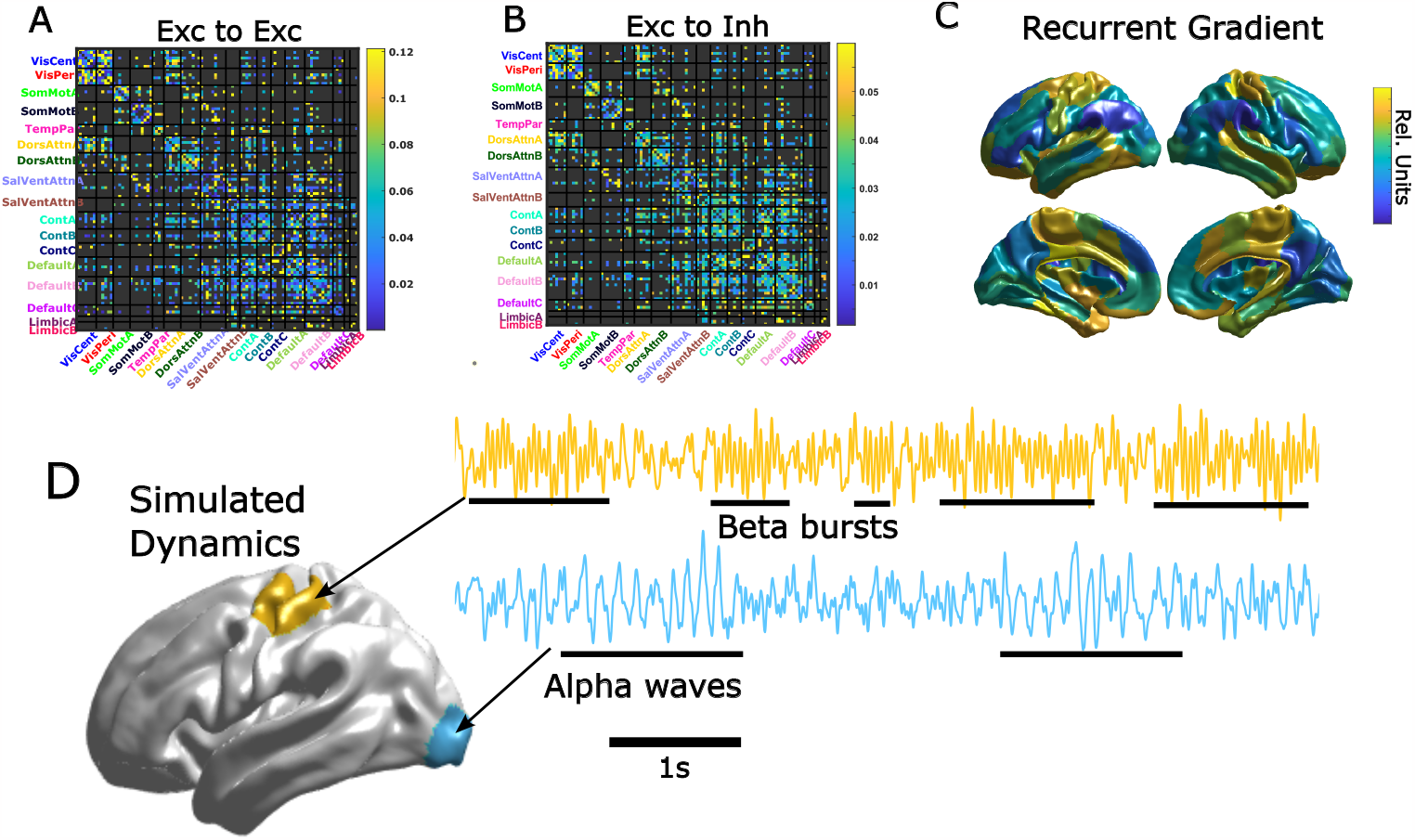
Group-average parameters and representative dynamics. A) Group-average excitatory-to-excitatory connection matrix sorted by the Yeo-17 networks [33]. Dark-grey connections denote those which are not admissible as determined by the connectivity mask (see Sec. 8.6). B) Same as A) but for excitatory-to-inhibitory connections. C) Group-average spatial gradient of recurrent connections. Note that the separate recurrent connection types (EE, EI, II, IE) are derived from affine transformations of this gradient (see Sec. 8.7). D) Model-simulated activity motor (top) and visual (bottom) parcels of a representative HCP MEG subject. We note that the simulated brain activity is highly non-stationary and spatially heterogeneous. We highlight the spontaneous generation of narrow-band bursts (beta and alpha, respectively) interspersed with wider-band oscillations.

### 2.3. Models recapitulate and explain well-known electrophysioligical oscillations

We next analyzed the model dynamics (see Fig. 3D for an example timeseries). We first tested our models’ ability to correctly replicate the anatomical distributions of spectral power within the data (we refer to these as “spatiospectral features” for brevity). We divided the spectrum as follows: delta (1.5-4 Hz), theta (4-8 Hz), alpha (8-15 Hz), low beta (15-26 Hz), high beta (26-35 Hz), and gamma (*>*35 Hz) in accordance with the HCP MEG pipelines [25]. We normalized spectral power to have a sum of one across bands, within subject, for both data and models. In all spatial comparisons we use the precalculated source-level empirical estimates provided with the HCP ICA-MNE pipeline. We note that these analyses are not direct ground-truth tests, since the source-level empirical estimates are themselves limited in spatial resolution and likely overestimate smoothness. Hence, we do not expect exact agreement on parcel-level values, particularly near the boundaries between brain networks. At these boundaries, models produce much sharper spatial divisions than the empirical source estimates (see e.g., Fig. 4 A,B). We pay special attention to the alpha and beta spectral bands as these are most prominent in resting-state.

**Figure 4.**
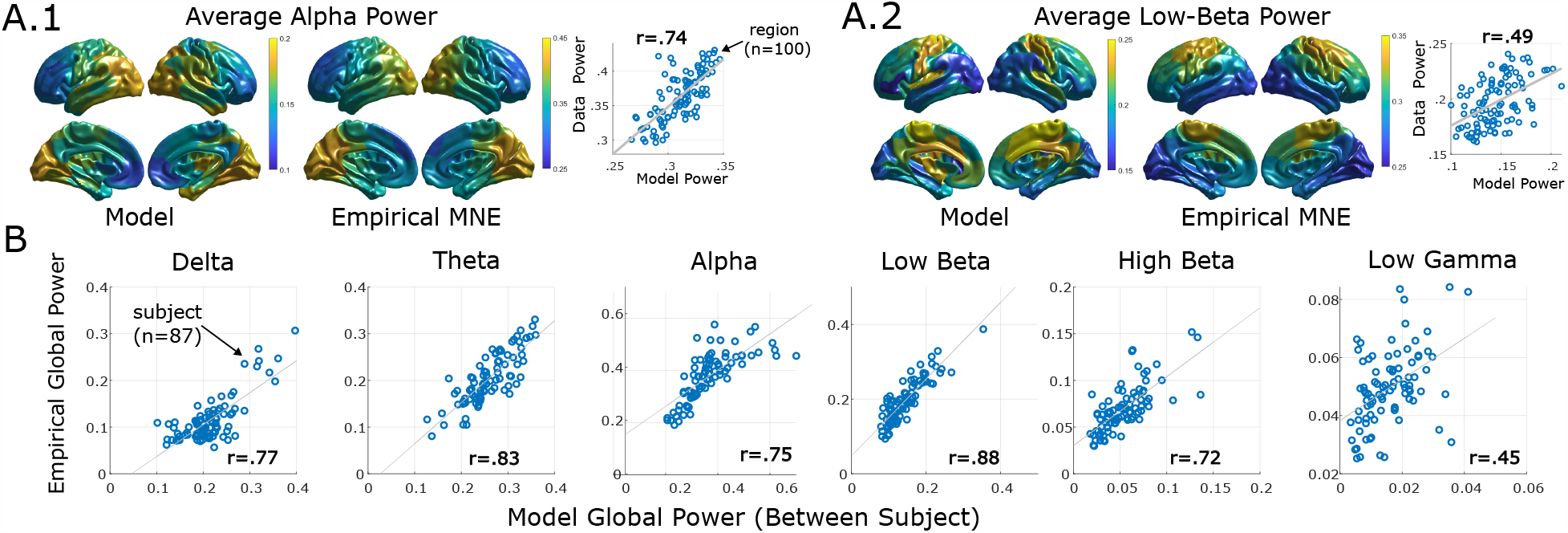
Precision brain models replicate empirical spatiospectral patterns. A) Group-level anatomical distributions of spectral power strongly correlate between model-predictions and source-localized MEG for two of the stereotyped M/EEG bands at rest: alpha (A.1) and low-beta (A.2). B) Individual differences in whole-brain spectral power (averaged over parcels) are reproduced across frequency bands by individiualized brain models.

We first analyzed results at group level by comparing the group-average anatomical-profile of spectral power between model simulations and empirical estimates. Results demonstrate good spatial agreement for the alpha band (*r*(98) = .74; Fig. 4A.1) and moderate agreement for the beta bands (*r*(98) = .49, *r*(98) = .42, respectively; Fig. 4A.2). We note that the model-predicted low-beta is highly localized to the somatomotor network, compared to the blurrier source-estimates. The slow delta band also exhibited high similarity between data and models (*r*(98) = .64). By contrast, the theta band was moderately consistent: *r*(98) = .52.

We next examined model fidelity in replicating oscillatory dynamics at the individual level. At the coarsest level, we found that models strongly reproduce individual differences in global spectral power (whole-brain average) across spectral bands in the training data (delta through high-beta: *p*^′^ *s <* 2.2E-15, Fig. 4B). Interestingly, we also found a moderate correlation between individual differences in the low-gamma band (*r*(85) = .45, *p <*1.1E-5, Fig. 4B) despite its suppression in training-data due to 30Hz low-pass filtering (excluding the full band). However, in contrast to other bands, the magnitude of model-predicted low-gamma power was significantly smaller than the HCP source-estimates and the average spatial profile was not consistent with data (*r*(98) = .07), so this result may simply reflect residual gamma-power retained in training data even after filtering.

We also found that models replicated individual differences in spectral power at the network-level for the main resting-state bands (alpha, low-beta). To compute network-level power, we averaged spectral power among parcels belonging to each of the 17 Yeo [33] resting-state networks as implemented in the Schaefer 100-17network parcellation [35]. We correlated model-predicted and empirical power in a multilevel model with a fixed-effect of subject (global power) collapsed across all networks. We found the strongest similarity between model-predictions and data for the alpha (*r*(1390) = .54), low-beta (*r* = .50) and delta (*r* = .43) bands. While all statistical models were significant (due to the large number of data points), similarity was weak in the theta (*r* = .35), high-beta (*r* = .32), and gamma (*r* = .10) bands. These results suggest that, for the main resting-state bands (alpha, low-beta), models correctly predict network-level power at the single subject-level. However, despite high accuracy in predicting global power (see above), models are less accurate at predicting the anatomical/network sources of high-beta and theta-band power.

As a final validation, we examined whether models predict individual variation in the peak-frequency of the alpha band. Empirically, this characterization has proven a remarkably stable and predictive measure of individual differences in brain and behavior [4, 36]. The functional significance of “peak-alpha” is not yet resolved, although many accounts posit that the width of an alpha-oscillation determines the temporal window over which phase-linked processes (e.g., information integration) occur [37, 5]. We found that models were accurate at reproducing the global individual alpha-frequencies (*r*(85) = .625, *p* ≈ 0; Fig. 5A). At the parcel-level, we found that model accuracy was highest in predicting peak-alpha within posterior cortex which agrees with the anatomical expression of alpha power (Fig. 5B).

**Figure 5.**
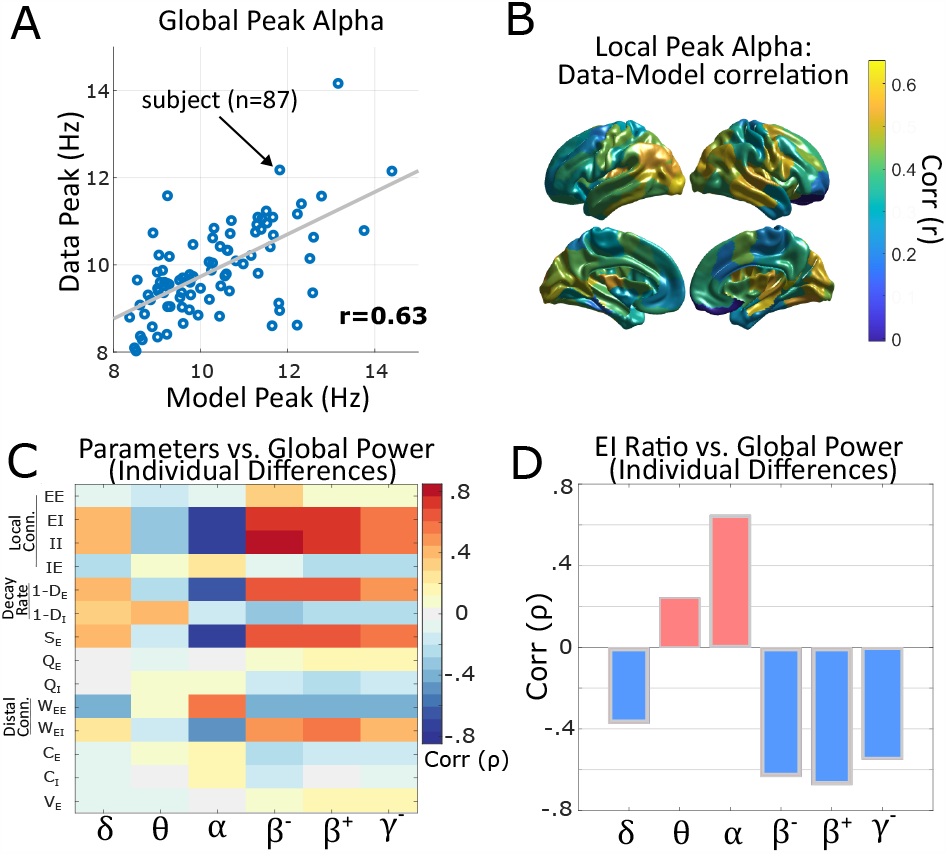
Models predict individual’s peak frequency within the alpha band and explain individual differences in power. A) Correlation between predicted and empirical global-peak alpha (average over parcels). B) Model predictions of local peak-frequency are most accurate in posterior regions in which the alpha rhythm is dominant. C) Spearman correlation matrix between model parameters (see Sec 8.1 for definitions) and global power in each frequency band. D) Correlations between individual differences in the ratio of excitatory and inhibitory activity and global power by frequency band.

### 2.4. E-I balance predicts individual differences in whole-brain average spectral power

Generative models, as we present here, can form predictions using either overt linkages to model-parameters or as emergent phenomena generated by their dynamics. We therefore tested whether individual differences in spectral power or peak-alpha are correlated with individual parameters embedded in the models (Fig. 5C). Between subjects, we found that connections onto inhibitory populations (local *EI*, local *II*, and distal *EI*) had a strong negative correlation with alpha-power (*ρ*(85) = −.73, −.75, −.51; *p*’s*<*E-6, respectively), and a positive correlation with beta-power (low beta: *ρ* = .74, .78, .48; *p*’s*<*3E-6, high-beta: *ρ* = .72, .76, .67; *p*’s*<*2E-8). This relationship also held for the excitatory decay-rate (alpha: *ρ* = −.64, low-beta: *ρ* = .64, high-beta: *ρ* = .64; *p*’s*<*E-10) and was reversed for distal *EE* connections (alpha, *ρ* = .54, low-beta: *ρ* = −.43, high-beta: *ρ* = −.42, *p*’s*<*6E-4). Relationships were similar to alpha, but weaker, in the theta band and similar to beta in the low-gamma band (Fig. 5C). These parameter-level relationships suggest that the global (i.e., whole-brain average) excitation-inhibition ratio changes spectral power in the low vs. high frequency bands.

Motivated by these parameter-level relationships, we quantified the model-predicted excitation-inhibition ratio for each parcel using the standard-deviation of simulated timeseries: *σ*(*x*^*E*^)*/σ*(*x*^*I*^). This estimate was highly reliable (for brain-wide average: *ICC* = .78) with much larger variation between-subject (*σ* = 1.46) than between-region (*σ* = .10), hence we only further investigated the brain-wide average due to small anatomical variation. In agreement with the aforementioned parameters, the predicted EI-ratio was positively correlated with global alpha-power (*ρ* = .65, *p* ≈ 0, Fig. 5D), but negatively with beta power (low-beta: *ρ* = −.63, high-beta: *ρ* = −.67, *p*’s≈ 0). Results thus indicate that individual differences in models’ excitation-inhibition ratio predict empirical power in the higher-frequency bands. This result is complementary, but not identical, to theoretical models suggesting a relationship between excitation-inhibition ratio and the 1*/ f* slope of power-spectral density [38, 39]. Interestingly, however, these relationships only held at the global scale. Neither the model-predicted excitation-inhibition ratio, nor the strength of distal connections (EE, EI) predicted the anatomical distribution of spectral power for any band (max *ρ*(98) = .24, n.s.). The spatial gradient of local-recurrent connections was weakly correlated with gamma-band power *ρ*(98) = .31, *p* = .040 post-Bonferroni correction) and non-significant for all other bands. We did not find any significant relationships between model parameters/excitation-inhibition ratio and peak alpha frequency in terms of individual differences or anatomy.

Thus, in total, individual model parameters strongly predict individual differences in global beta vs. alpha power, via the excitation-inhibition ratio. But, these parameters do not predict the *anatomical* profile of spectral power or the peak alpha frequency, even though these features *are* readily generated by the models (see Sec. 2.3 above). Thus, the spatial distribution of spectral power is an emergent property generated by model dynamics as opposed to reflecting a singular of the model.

### 2.5. Model predicts the existence of individually reliable equilibrium and non-equilibrium oscillatory dynamics

Lastly, we used the models to interrogate the dynamical properties of two resting-state oscillations: alpha waves and beta waves (low-beta and high-beta). As previously mentioned, alpha rhythms (defined as 8-15Hz for HCP, but more commonly 8-12Hz) are high-amplitude posterior oscillations, often present at rest, which are amplified during eye closure and by tasks that require visual inhibition, broadly defined. While alpha constitutes one of the first discovered waking-EEG components, theories regarding its origin and mechanism have evolved significantly over the past two decades. Initial theories, based upon the eye-closure effect, posited that alpha represented a default “cortical idling” state to which the visual system would return, absent environmental stimuli (see [40, 41] for review), potentially driven by thalamic nuclei [1, 2]. By contrast, alpha activity is now largely interpreted through the lens of preparatory attention [42] and active visual inhibition (e.g. [43, 44]). Several recent results have also indicated the potential for cortically-initiated alpha waves to propagate retrograde along the dorsal visual pathway, from anterior (higher-order) to posterior (lower-order) cortex [3, 45], in addition to a visually-evoked forward-propagating wave [45, 46].

We tested dynamical mechanisms by which alpha and beta waves could be generated at rest. From a theoretical perspective, there are several candidate mechanisms that can generate wave-like dynamics, including: limit cycles (stable, periodic behavior that the system will recover after small perturbations), quasiperiodic behavior (mixed oscillations at incommensurable frequencies), aperiodic waves (e.g. spectrally-concentrated chaos), stable foci/spiral-points (transient damped-oscillations when the system is perturbed), and noise-driven ‘switching’ between different equilibrium states. As an initial investigation we considered the attractor structure of the models, dividing models into those with equilibrium-style attractors and those with non-equilibrium attractors. The former class defines models which, in the absence of external perturbation, generate complex transient behavior, but will eventually settle into a steady-state. The latter class of dynamics, converge onto low-dimensional patterns of persistent activity, such as oscillations, even without perturbation. The dynamics embedded near an attractor thus determine the system’s “default” mode of activity, whereas transient patterns of activity require some initial perturbation into that regime.

We found that these categories were consistent within-subject with 78% of subjects having the same category for separate models fit to each data half (Odds Ratio=14.4, independence *χ*^2^(1) = 27.8, *p <* 1.4*E* − 7). Of the 87 subjects, 40 (46%) belonged to the equilibrium-group for both scans, 28 (32%) belonged to the non-equilibrium group, and 19 (22%) had one scan in each category (see Fig. 6A). Surprisingly, these categories proved powerful markers of individual differences in empirical spectral power. We observed strong increases in alpha and theta power, but decreases in beta and low-gamma power for subjects with non-equilibrium dynamics (theta: *t*(66) = 3.54, alpha: *t* = 6.1, low-beta: *t* = −8.57, high-beta: *t* = −6.80, low-gamma: *t* = −6.48, *p*’s*<*.0008) (Fig. 6B). In a agreement with the group-wide parameter-correlation results (Fig. 5), we found that equilibrium (low-alpha) subjects had an increase in EI connection strength leading to a lower ratio of excitatory-to-inhibitory activity (Fig. 6C) and changes in the integration time-constants. As before, we found contrasting influences of the *II* strength and inhibitory decay-parameter, however the net influence, over relevant ranges of activity, was an increase in negative feedback (faster decay) for the equilibrium subjects, particularly near the equilibrium (where *ψ*’ is particularly large). However, there was no difference between groups for the empirical peak-alpha frequency (*t*(66) = −1.83, n.s.). These results suggest that some aspect of the low-dimensional attractors promote the generation of high-amplitude oscillations in a band-selective manner as opposed to globally speeding/slowing dynamics (which would be reflected in peak frequency).

**Figure 6.**
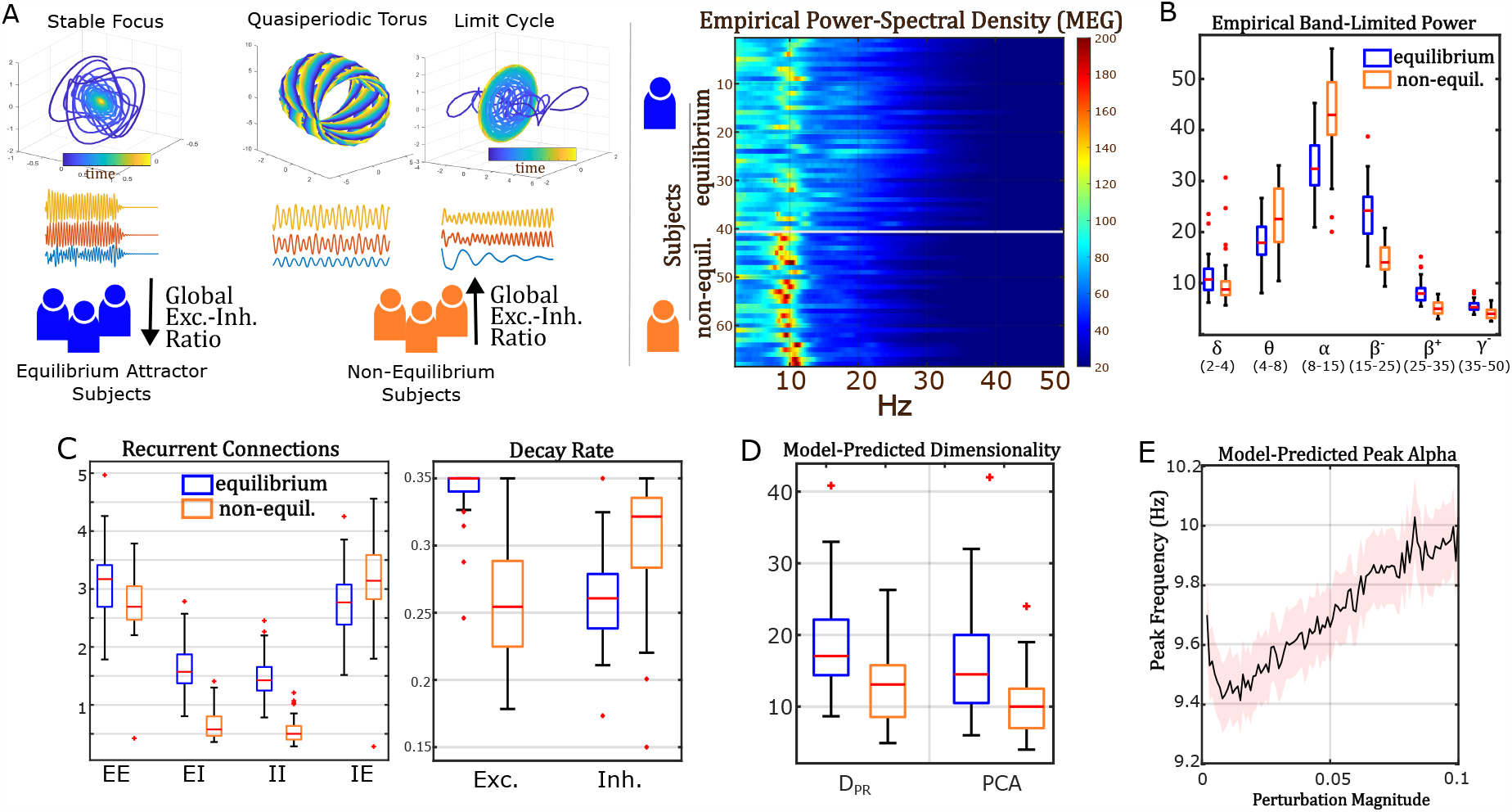
Models identify a taxonomy of subjects based upon attractor geometry. A) Global power-spectral density for all subjects in which models were consistent in producing equilibrium or non-equilibrium dynamics. Left side shows example attractors for three subjects, one with an equilibrium spiral-point attractor (top) and two with non-equilibrium attractors (a quasiperiodic torus on the left and limit-cycle on the right). All attractors are projected onto 3-dimensional coordinates identified by PCA. Right side shows brainwide average power-spectral density by subject. B) Band limited power for each group of subjects. C) Models predict that attractor geometry is associated with alterations in local excitatory-inhibitory balance and timescales. D) “Dimensionality” of model dynamics by attractor group as calculated using either Participation-Ratio Dimension or thresholded-PCA (Sec. 4.6). E) Models predict that alpha oscillations become faster (higher-peak frequency) under perturbation. Shading indicates ±1 standard-error (n=87).

### 2.6. Alpha oscillations arise from low-dimensional dynamics and are sensitive to perturbation

As an additional test, we considered how the induced dimension of model dynamics affect the alpha rhythm. Here, we are interested in the effective dimension of the stochastic (noisy) dynamics, i.e., how expressive are the models in terms of spatial activity patterns under physiological conditions. For this test, we used two different component-based measures of dimensionality for comparison: a hard-threshold dimension based upon the number of nontrivial PCA components (*D*_*PCA*_, threshold = .1*λ*_*max*_) and the Participation Ratio Dimension (*D*_*PR*_, see 4.6) which is a graded measure. We did not use topological definitions (e.g. Hausdorff dimension) as we were interested in global dynamics embedded within the high-dimensional space, as opposed to only studying the attractors. Results indicated lower induced-dimension for non-equilibrium subjects (*D*_*PR*_ = 16.3 ± 7.9, *D*_*PCA*_ = 10.4 ± 5.0) compared with equilibrium subjects (*D*_*PR*_ = 18.9 ± 7.4, *D*_*PCA*_ = 13.2 ± 5.3) as assessed with two-sample t-tests corrected for unequal variance (*D*_*PR*_ : *t*(65.9) = −3.7, *p <* .0005, *D*_*PCA*_ : *t*(65.5) = −3.8, *p <* .0004; Fig. 6D). These findings agree with previous, empirical descriptions of lower-dimensional dynamics associated with alpha band vs. beta ([47]) activity. This analysis indicates that the lower-dimensional dynamics which embed non-equilibrium attractors, contract dynamics throughout the state-space, rather than solely in the vicinity of the attractor. Thus, the global dynamics of non-equilibrium subjects are shaped by lower-dimensional structures.

To clarify the association of low-dimensional/ nonequilibrium dynamics with spectral power, we investigated how the models reacted to perturbations. This analysis is important for interrogating whether the alpha rhythm is, itself, an attractor (e.g., limit-cycle) or rather reflects transient behavior built upon the nearby dynamics. At the systems-level, the former case corresponds to a default-behavior which cannot be suppressed without external input (the historical alpha interpretation) whereas the latter represents a regime that leverages the intrinsic dynamics, but requires some perturbation to initialize. To differentiate these cases, we examined the model response under perturbations by simulating the models in the presence or absence of intrinsic noise. As previously indicated, noisy-simulations of non-equilibrium subjects predict greater alpha and lower beta power than equilibrium subjects in agreement with the data (see above, Fig. 6A,B). However, in the absence of intrinsic noise, model dynamics converge onto the attractors. We found that most non-equilibrium attractors had greatest power above 20Hz, despite the models generating lower beta-power in the presence of noise. When a spectral component in the alpha-range was present, the other frequencies were not harmonics of the alpha-component. These results suggest that the alpha oscillation builds off of low-dimensional intrinsic dynamics, but is not itself a self-sustaining behavior. This results indicate that, for eyes-open resting-state, active perturbations are required and sustain an alpha wave, although such perturbations need not be large nor applied continuously.

On this point, we examined how perturbations affect model dynamics. For this analysis, we focused upon changes within the alpha spectra, as opposed to between spectra. We also emphasize high-level properties, as opposed to simulating a specific task, so we modeled environmental perturbations as a random process independently delivered to each brain area. The perturbation magnitude is thus equivalent to the noise-level which we applied as a scaling-factor to simulated physiological/process noise (with baseline covariance *Q* as estimated by individual models). As before, process noise denotes stochasticity which drives a system, as opposed to artifact which only appears in measurements. We scaled noise-perturbations from 2% to 100% of the variance used in simulations, with a resolution of 1%. Averaging over subjects, we found that the peak alpha frequency within the visual network linearly increased as a function of perturbation strength (*r*(97) = .96, *p* ≈ 0; Fig. 6E). This effect agrees with empirical findings of the peak alpha frequency changing within-subject, depending upon the level of engagement (increases during task, [4, 5]) and has been explored as a generic property in previous theoretical models [48, 49]. We also note the observed change in peak-frequency indicates inherently nonlinear dynamics, so the existence of an equilibrium attractor should not be confused with approximately-linear dynamics. Together, these demonstrations indicate the potential of our framework to inform high-level systems neuroscience and implicate mechanisms of person-driven and context-driven variation.

## 3. Discussion

We have presented a novel framework for estimating precision brain models from single-subject M/EEG data. Our gBPKF algorithm is shown highly capable of solving the dual-estimation problems inherent in direct brain modeling and reliably estimates latent brain model parameters directly from M/EEG timeseries. Importantly, we stress that our approach directly applies nonlinear system-identification to macroscale cortical activity, seeking to solve for the system’s vector-field (i.e., the moment-to-moment variation) as opposed to replicating a specific signal feature. Nonetheless, our approach reproduces the spatial distributions and individual differences in band-limited power across the main M/EEG bands with high fidelity. The models also link individual differences in alpha power to the geometry of underlying dynamics as shaped by excitatory-inhibitory balance. These applications demonstrate the inferential power of individualized (precision) brain-modeling.

In the present work, we demonstrated our technique using magnetoencephalography (MEG) data from the Human Connectome Project (HCP; [25]). However, we expect that the approach will be similarly useful to estimating brain models from other fast-timescale modalities, particularly electroencephalography (EEG). These approaches also measure the effect of cortical dipoles at a distance and can be similarly modeled in a general state-space framework. One difference, however, is that voltage-based modalities such as EEG, ECoG etc., are inherently referential i.e., they measure the electrical potential between points. Fortunately, this aspect is fully compatible with our algorithm and simply corresponds to a different forward model/measurement matrix (*H*). On a technical note, care should be taken so that the resultant data is full-rank (e.g. by factoring *H*; see Sec 8.8) as the leadfield is guaranteed to be rank-deficient (i.e. *HH*^*T*^ is not invertible) when mean-referencing is applied. The referential nature of voltage also generates a new shift invariance (nonuniqueness of *C* and *V* values) although this effect only applies to their absolute values, as opposed to the relative spatial patterns. However, we believe that the general enterprise of characterizing individual brain dynamics and estimating models will prove similarly applicable to EEG as MEG.

We view our present model as having two primary limitations: 1) assumptions regarding how the M/EEG signal is generated and 2) the omission of subcortical sources. We stress that these limitations are features of our current model (two nodes per cortical parcel) rather than the approach *perse*. We have designed our publicly-available code so that it is easy to implement arbitrary models of neural circuitry and M/EEG signal weighting (with the usual trade-offs between run-time/robustness and model complexity). We therefore examine some of our model’s limitations with the caveat that these limitations are not inherent to our general estimation approach (gBPKF). In the first limitation, we make the typical assumption that current-dipoles reflect excitatory neuron depolarization and that the dipole, like the cells themselves, is oriented normal the cortical surface. While convenient, these assumptions are not always valid ([24, 50]) so further refinement of the measurement matrix (*H*) may improve anatomical precision.

A second limitation arises from the neglect of subcortical influences. In the current work, we chose to only model cortex, in line with the predominantly cortical origin of M/EEG signals. However, interactions with the brainstem, thalamus, and hippocampus are known to generate many slow EEG components. It is for this reason that our frequency-domain validations focused upon the faster alpha and beta bands, although the cortical vs. subcortical origins of either phenomena are not yet settled (although see [3]). In any case, it is clear that subcortical influences certainly guide cortical dynamics and that their inclusion could improve model performance. Several Dynamic Causal Modeling approaches, for instance, have included subcortical regions in modeling M/EEG ([51, 52]). This approach usually requires strong priors as the optimization may otherwise become ill-posed. By contrast, we have first prioritized validating our framework within an empirically-verifiable model space, as opposed to including as many degrees of freedom as possible. It is therefore possible that some of the would-be explanatory variance associated with cortico-thalamo-cortical pathways is mismodeled in our approach as direct corticocortical connections in the absence of a subcortical component. However, we have designed our modeling approach such that subcortical regions can be easily added with or without various priors as another latent-variable, treated analogously to the interneurons (*H*_*subcort*_ = 0). We encourage (verifiable) expansion in this direction.

In conclusion, we have validated a new approach for precision brain-modeling using single-subject M/EEG and illustrated its explanatory potential in the context of resting-state oscillations. We hope that these innovations will enable new insight into individual variation and circuit mechanisms that may, eventually, inform new ways of interacting with the brain.

## 4. Data Processing and Statistical Methods

In this section we briefly describe the resting-state magnetoencephalography (MEG) data used for model training and validation. An introduction to the Kalman Filter and details of the gBPKF algorithm are presented in the SI. Technical descriptions of ground-truth model generation are also provided in the SI.

### 4.1. HCP MEG Data Processing

Models were fit to magnetoencephalography (MEG) data provided by the Human Connectome Project (HCP; [25]). We used the minimally-preprocessed HCP pipeline for MEG which centers on using Independent Component Analysis (ICA) to identify and remove artifact. Thus we used the HCP MEG data as-provided up to the ICA-removal step. Whereas the HCP pipeline proceeds using only the “good” independent components (IC’s), we instead projected-out the bad ICs from the sensor-level timeseries. While both approaches remove the “bad” IC’s as identified in the HCP MEG release, our approach still retains the original sensor space and the associated low-variance IC’s which the HCP considered neglible. The rationale for this deviation is so that the those dimensions of measurement are retained as the absence of (significant) signal along those dimensions is itself informative.

We then projected data according to the left singular-vectors of the leadfield matrix (derived from the pre-calculated Boundary Element Method headmodels). This step corresponds to reducing dimensionality based upon what spatial patterns MEG *can* measure as opposed to those that were observed (in practice these approaches overlap). Our criterion was to only retain leadfield dimensions (singular vectors whose singular values were ≥ 1% of the maximum singular value. These reductions were done on a per-scan basis which meant that, due to the removal of bad channels, projections (hence measurement models) often differed between scans of the same subject.

### 4.2. Frequency-Domain Filtering

Data was filtered between the delta and high-beta bands (1.3-30Hz). The 30Hz upper limit marks the high-beta band while the 1.3Hz lower limit was adopted from the HCP ICA processing pipeline and does not include the full delta-band (i.e., is above the slowest delta waves). This range also includes the full alpha and theta-bands. As part of the HCP MEG pipeline, high-artifact data segments are automatically removed, resulting in variable length segments of good-quality data. To calculate power spectral density (PSD) at the subject-level we first discarded segments lasting less than 20s. The PSD was then calculated for each timeseries and discretized with resolution 0.25 Hz. The resultant PSDs were averaged across segments, weighted according to segment length.

For band-limited power, we used the pre-calculated HCP estimates in the ICA-MNE pipeline. This pipeline solves for a full dipole-vector timeseries at each of the 8004 vertices (3 dipole coordinates per vertex). Band-limited power is estimated by filtering the timeseries for each dipole coordinate and then calculating the squared magnitude of the filtered dipole vectors. HCP defines 8 specific bands: delta, theta, alpha, low-beta, high-beta, and low/mid/high gamma. In the presented results, we normalized the average band-limited power at each dipole by the whole-spectrum (unfiltered) power and then averaged within parcel to get parcel-level power.

### 4.3. ICA Artifact Detection

We used the HCP MEG-ICA pipeline to identify artifactual Independent Components (ICs). The HCP MEG2 and later releases sorts large ICs as having a brain or non-brain origin. However, whereas the original ICA pipeline retained only “brain” IC’s (removing both artifactual and low-variance IC’s) we simply removed the non-brain IC’s by orthogonal projection, thereby retaining the original channel dimensions. This step retains information that the low-variance IC’s are measurable and small, whereas removing them would imply that those dimensions are unmeasurable (potentially leading to model overfit). Denoting the ICA mixing matrix for the bad ICs as *M*_*bad*_ and the post-ICA corrected measurements *y*^*ICA*^:

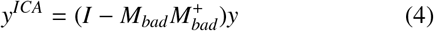

### 4.4. Source Localization

We used the standard HCP anatomical pipeline to compute forward head models. Comparisons with empirical source-level power all used the HCP pre-calculated power-distributions which allow full 3d dipole configurations ([25]). However, model-training requires a fixed mapping between activity patterns and measurements, and thus a single, constant, dipole orientation per vertex (up to sign reversal). We reduced the 3d forward model to a single dipole direction per vertex by assuming that dipoles are oriented normal to the cortical surface (as calculated in FieldTrip using the vertex cross-product method).

In order to transform MEG magnetometer/gradiometer measurements onto cortical dipoles we used Minimum Norm Estimation (MNE; [53]). Like other linear source-localizers, The MNE inverse matrix ℳ maps sensor-level data onto the brain:

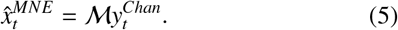

Conceptually, MNE minimizes the expected mean-square-error (like the Kalman Filter) and therefore takes the regression form:

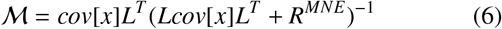

Following the HCP ICA-MNE pipeline, we use the simplified assumption that *cov*[*x*] and *R*^*MNE*^ are each independent and identically-distributed (iid) hence, using the noise-signal-ratio (NSR) coefficient *λ*:

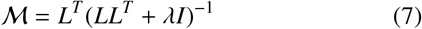

We calculated *λ* analogous to the HCP minimal pipeline, but with a single *λ* per run, applied in channel-space, whereas the HCP rescaled *λ*’s for each IC. This difference is because we retained the original channel space, instead of reducing dimensionality to the IC space.

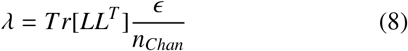

with noise-value factor *ϵ* = 8. The inverse solution ℳ was calculated separately for each resting-state run and potentially differed (e.g. do to noisy channel removal). We further rescaled each row of ℳ to produce unit variance in 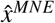 which was done separately for each run.

### 4.5. Statistical Analysis

Our analyses generally fall into three categories: 1) validating/benchmarking the approach with a known (simulated) ground-truth, 2) assessing reliability with real-world data, and 3) exploring dynamical predictions made by the models. For these analyses we used three similarity measures depending upon the variable’s dimensionality. We used simple correlation (collapsing over non-masked connections) to gauge the overall similarity of two matrices/vectors. To assess the reliability of individual differences, we used Intra-Class Correlation ([54]) for scalar-valued parameters (e.g., time constants) and Image Intra-Class Correlation (I2C2; [34]) for multivariate parameters (e.g. connectivity). All reported *p*-values are 2-tailed. Multiple-comparison corrections all used the Bonferroni method and statistics reported as significant all passed this threshold. We use the notation *p* ≈ 0 for calculated *p* values less than 10^−10^ as exact estimates are likely inaccurate past this point. When multiple related analyses are presented in-text, we typically report the largest *p*-value over all of the analyses with the notation *p*^′^ *s <*, to improve readability.

### 4.6. Quantifying Dimensionality

We quantified dimensionality of stochastic dynamics in two ways, both based off of the covariance eigenspectrum, with convergent results. First, we used the PCA-threshold method with a hard-boundary defined by 1% of the maximal component weight (eigenvalue). For this method, we quantified dimensionality as the number of components passing this threshold. For comparison, we used the participation ratio dimension (*D*_*PR*_, [55, 56]) which is calculated from the covariance eigenvalues (*λ*):

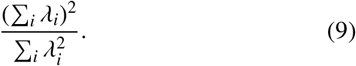

We used *D*_*PR*_ for comparison as it provides a soft dimensionality metric in terms of the variance spread as opposed to being premised upon a single, low-dimensional surface. It is also sensitive to dynamics which are nontrivial in the stochastic case, but eventually converge in the absence of noise. We present *D*_*PR*_ calculated using z-scored simulated data (i.e., using correlation instead of covariance) to control for the different scaling of excitatory and inhibitory neurons. However, statistical inferences led to the same conclusion with/without normalization.

## 5. Data and Code Availability

Resting-state MEG data is publicly available through the Human Connectome Project (HCP; [25]). It can be accessed, through a registered account, at db.humanconnectome.org. Data processing code, as described below, is available through HCP. Interested users should download the “megconnectome” pipeline scripts through the HCP database. A software package containing MATLAB code for gBPKF model-fitting, simulation, and visualizing results is available at the primary author’s github.

## 6. Author Contributions

M.S. designed and performed analyses, acquired funding, and wrote the paper. T.B., M.C., and S.C. wrote the paper and supervised the study.

## 7. Acknowledgements

MS was funded by NSF-DGE-1143954 from the US National Science Foundation, the McDonnell Center for Systems Neuroscience and NIH T32 DA007261-29 from the National Institute on Drug Addiction. Portions of this work were supported by NSF 1653589 and NSF 1835209 (SC), from the US National Science Foundation and NIMH Administrative Supplement MH066078-15S1 (TB).

## 8. Supplementary Information

### 8.1. Mesoscale Individualized NeuroDynamic Modeling at Fast Timescales

We first describe the generative model to be estimated by each subject’s data. Our model formulation is similar in motivation to conventional neural mass models (e.g., [26]). Each brain region contains two neural populations: an excitatory and an inhibitory population. A sigmoidal nonlinearity (*ψ*) converts population-average activation (an abstraction of depolarization) into a normalized output (analogous to firing rate). The shape of the nonlinear function is parameterized by a set of unknown variables (*α*). Macroscale electromagnetic fields (MEG, EEG, LFP, etc.) generated by the brain derive primarily from post-synaptic and dendritic potentials as opposed to action-potentials, hence, for our data-driven models, we use activation/depolarization (as opposed to firing-rate) as the state variable, denoted (*x*).

Each population receives some baseline level of drive *c* and it returns to baseline at a rate (1-*D*) with *D* a diagonal matrix of autoregressive coefficients. We refer to the quantity (1-*D*) as the “decay” rate. Both excitatory and inhibitory cells connect locally and excitatory cells also connect to distal brain areas via connection matrices (*W*). A major difference in our model, from previous approaches, is that we consider two types of inter-regional connections from excitatory cells: excitatory-excitatory and excitatory-inhibitory, whereas previous approaches have been constrained, by the nature of diffusion data, to a single, undirected form of connectivity (e.g., [17]). The noise terms *ε* correspond to unmodeled physiological processes and are assumed to be Gaussian with zero mean and covariance *Q*_*t*_. Together these equations are:

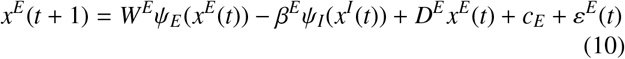

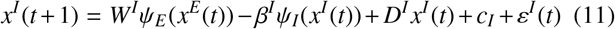

The nonlinear function *ψ* is specified analogous a 2-parameter logistic function with gains (*s*^*E*^,*s*^*I*^) and bias terms *v*^*E*^, *v*^*I*^. Here, the gain *s* is scalar-valued (independent of parcel), whereas the biases *v* vary by parcel. For simplicity, we used tanh for the nonlinear function and note that such models can be directly rewritten with nonnegative activation (logistic sigmoid), if desired. Without loss of generality (see Sec. 8.7) we fix *s*^*I*^ = 1, *v*^*I*^ = 0, so these parameters are only solved for the excitatory population. We condense the separate population equations into the more general form with excitatory and inhibitory activity concatenated as *x*_*t*_:

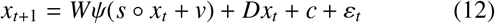

This is referred to as the state equation as it defines how the state-variable “x” evolves in time. Coupled with the state-equation is an associated measurement-equation (“forward model”) which defines how patterns of brain activation (*x*_*t*_) are reflected in sensor readings (*y*_*t*_). As we are interested in electromagnetic fields (which add), this transformation constitutes a linear mixing defined by the matrix *H* and sensor-level noise *η* (which evolves as a Gaussian process with zero-mean and covariance *R*_*t*_).

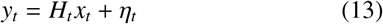

Previous research strongly indicates that brain potentials measured from the scalp (MEG and EEG) are primarily generated by cortical pyramidal cells, whose asymmetric geometry supports the formation of dipoles ([24]). By contrast, the symmetric geometry of inhibitory neurons (e.g., stellate cells) leads to the microscopic (subcellular) dipoles largely canceling when measured from a distance. Thus, for present purposes, we model the signal mixing matrix *H*_*t*_ as consisting of a forward matrix 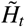 for excitatory populations (described in detail later) and zero for inhibitory populations. Similarly, we assume that dipoles are oriented normal to the cortical surface reflecting the underlying orientation of pyramidal cells. However, our methodology is relevant to any specification of measurement model and can therefore be adapted to alternative models of EEG signal generation (i.e., by defining a different 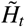 matrix). The measurement matrix *H* is allowed to be time-varying (e.g., for dropping a channel during periods of artifact).

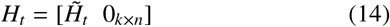

We assume that the following pieces of information are known (or well-approximated):

1. A forward measurement model (*H*_*t*_)
2. The sensor-noise covariance (*R*_*t*_)
3. Some reasonable restrictions on the connectivity graph

The stringency of the last requirement depends upon the dimensionality of the measurements (e.g., channel count) relative the number of brain areas, particularly those with low-SNR. In our case, we directly inferred all these properties from data by using MRI data to calculate forward models (boundary element method) and empty-room recordings to estimate *R*. We used the group-level distribution of fMRI MINDy models ([19]) to generate a binary “connectivity mask” of plausible connections (see Sec. 8.6). We used the same connectivity mask for excitatory-excitatory and excitatory-inhibitory connectomes. The remaining challenge consists of solving for the latent brain activity (*x*_*t*_) and brain model parameters (*Q, W, D, c, s, v*) given measurements *y*_*t*_. This endeavor is non-trivial as it involves a high-dimensional nonlinear optimization problem in which the system-states (brain activity for each population) are not directly accessible.

### 8.2. Kalman Filtering

The Kalman Filter is a recursive Bayesian algorithm for estimating *unknown* states of a *known* dynamical system. Given model parameters and measurements, the Kalman Filter and its nonlinear extensions, seek to minimize the expected difference (sum-of-squares) between true and estimated system states. The Kalman Filter differs from static approaches, however, in that its estimates also incorporate previous measurements.

At each time-step the Kalman Filter passes the estimated distributions through a noisy dynamical-systems model. Using the measurement model, the Kalman Filter updates this distribution based upon the new data. By tracking the distribution over time, the Kalman Filter detects latent state variables (e.g., inhibitory neuron activity) through their predicted influence on measured variables.

#### 8.2.1. The Prediction Step

The Kalman Filter consists of alternating prediction and correction steps. During the *Prediction step*, the dynamical systems model is used to propagate the current state distribution 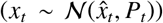 into an a-priori distribution for the next time step

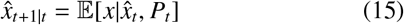

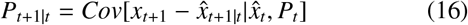

It is important to note that the covariance matrix here, (*P*_*t*_), is the error-covariance of estimating the true-state *x*_*t*_ using 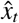 and not the covariance of a sample (i.e., *P*_*t*_ tracks uncertainty in estimating the vector *x*_*t*_ not the distribution of multiple states). The original Kalman Filter dealt with the linear case: *x*_*t*+1_ = *A*_*t*_ *x*_*t*_ + *ω*_*t*_ in which the solutions are:

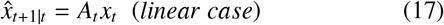

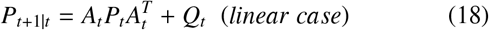

Different methods have been developed to extend this prediction step to the nonlinear case ([28, 29]), which we discuss later (Sec. 8.5).

#### 8.2.2. The Correction Step

The second step of the Kalman Filter uses new measurements to correct the a-priori predictions. This stage involves computing the Kalman Innovation (a prediction error) and adjusting state estimates proportionally to this error. Given prediction 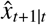 and new measurement *y*_*t*+1_, the prediction error (“Kalman Innovation”) is:

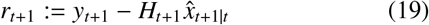

The Kalman correction occurs by multiplying this error by the Kalman-gain matrix (*K*_*t*+1_) to form an updated state estimate:

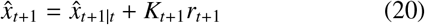

The gain, *K*_*t*+1_ is selected to minimize uncertainty (variance) in the corrected estimate:

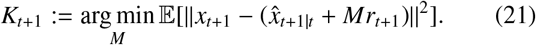

This problem constitutes least-squares regression and is therefore solved by:

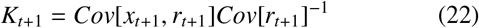

In our case, the model is linear in terms of measurements and measurement-noise (i.e. the M/EEG signal is a linear function of dipole strength), so the analytic solution is given by:

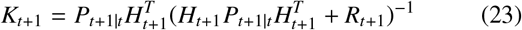

The posterior estimates, incorporating measurement *y*_*t*+1_ are thus:

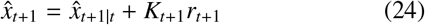

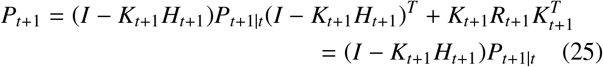

Following this correction, the Kalman Filter proceeds to the next time-step. Predictions, measurements and corrections are then assessed for *t* + 2. The major drawback of Kalman Filtering, however, is that it requires a known dynamical-systems model and initial distributions. In our framework, the dynamical-systems model is defined by the unknown parameters, so we express the Kalman Filter predictions as a function of these parameters which we optimize using gradient-based methods.

### 8.3. Optimization Objective

Our optimization framework in the current work solely seeks to minimize the error in predicting future sensor-level measurements, although associated code also enables the use of parameter-regularization and penalties based upon long-term model statistics (e.g. matching the observed covariance). For the present purpose, however, the model error corresponds to prediction error in the Kalman Filtering stage and in the free-running (forecasting) phase. For *k* Kalman-Filtering steps and *n* free-running prediction steps, we denote the error over start-times *t*_0_, we use 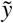 and 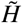 to indicate that comparison measurements used to evaluate error may be in a different space than those used to estimate states with the Kalman Filter:

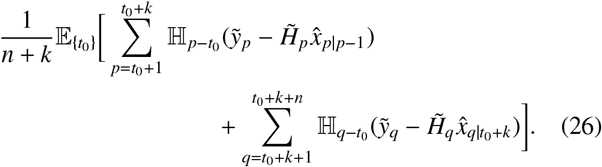

Thus, our cost function seeks to minimize the combined prediction error over both the Kalman-Filtering (left hand side) and forecasting (right hand side) steps. These errors are also averaged over all of the initial time points *t*_0_ (minibatch seed) which are randomly selected during each training iteration (see Sec. 8.9). Within our cost function, the term ℍ denotes the smooth Huber-loss transformation ([57, 58]):

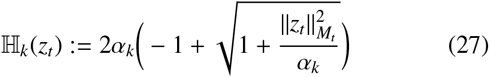

We applied the Huber-loss on-top of a conventional quadratic loss function with cost matrix *M*_*t*_ (see Sec. 8.8 for the choice of *M*):

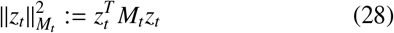

The smoothed Huber-loss function (ℍ) allows small values of ∥*z*∥^2^ to pass through unaltered, whereas large values of ∥*z*∥^2^ are quashed and limit to 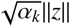. This smooth loss-function approximates the original Huber-loss which penalizes squared errors below a threshold and absolute errors above it. The Huber-loss is thus robust to outliers and improves model training in the presence of unusual events (e.g., artifact). The parameter *α*_*t*_ sets the soft-threshold for values to be quashed under Huber-loss. For each filtering/prediction step we set *α*_*k*_ equal to twice the expected median error for that step (i.e., *α*_1_ used the median of 1-step errors etc.) based upon previous iterations. This value was updated autoregressively after each training-batch to track the evolving error distribution (i.e., *α* became smaller as the model became more accurate). Denoting median by *med* and training iteration *m, α* was updated each training iteration according to:

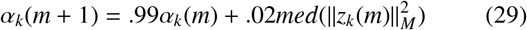

with the median being taken over initial conditions within a training minibatch. Note that this value is specific to the iteration within the prediction sequence (i.e., the *k*^*th*^-step prediction), not the recording time, hence ℍ is indexed by *p* − *t*_0_ and *q* − *t*_0_ in the cost function.

### 8.4. The generalized Back-Propagated Kalman Filter (gBPKF)

We first motivate our algorithm conceptually, by considering the dual-estimation of states and parameters as a min-min optimization. Namely, suppose that we have some cost-function *J* indicating the goodness-of-fit between model predictions (ŷ) and recorded data (*y*) by penalizing their difference. Because model predictions are a function of the system’s current state (*x*) the cost-function takes the form:

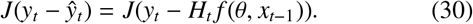

In the traditional model-fitting case, one solves for *θ* using *x* and *y*. However, M/EEG do not directly measure cellular activity, so both *x* and *θ* are unknown. A naive approach would be to directly optimize over both 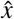 and *θ*; yet such an approach quickly explodes in the number of unknown variables. However, an alternative is to replace *x*_*t*−1_ with its best possible estimator, given the data. This estimate is a function of both the previous measurements (*y*_{*k<t*}_) and the system’s dynamics (as modeled by parameters *θ*). Since the true values are unknown, we define this estimator in terms of expected error:

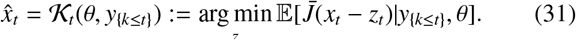

Here, 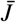 denotes whatever criterion is used to define a “good” state-estimate which need not be the same as the cost-function for optimizing the model (*J*). Substituting for 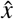 in the previous equation, gives an equation solely in terms of the unknown parameters (*θ*) and the measurements recorded up to time *t*:

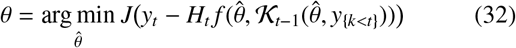

In our case, 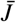 corresponds to sum-of-squares differences, hence the state estimate 𝒦 becomes the Kalman Filter (see Sec. 8.2,[59]). However, two elements of the Kalman Filter remain unknown: the initial state mean and covariance. To estimate these aspects we simulate the current model at the start of each iteration to derive baseline state (*x*_*t*_) distributions. This procedure constitutes the ”generative stage”. These distributions then parameterize a static linear filter (total-least squares) which forms the initial state expectation and error-covariance. Unlike source reconstruction, these initial estimates include both excitatory and inhibitory cells. This difference is because the simulations provide the steady-state covariance between all populations (excitatory and inhibitory).

Denoting the concatenated free-running simulation data *X*_*sim*_ and initial measurement *y*_0_ the initial state estimate and covariance is given by:

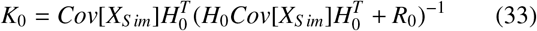

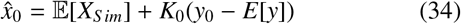

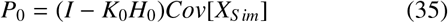

Note that these equations are identical to the Kalman update step (Eqs. 20,23) and, under trivializing assumptions, reduce to Minimum Norm Estimation (MNE; [53]) as is used in conventional source-localization for M/EEG. The initial state distribution parameters (expectation and covariance) are fed into a nonlinear Kalman Filtering algorithm. The Kalman Filter integrates a sequence of “k” measurement timesteps to refined estimates of latent neural activity. We then use the last state-estimate provided by the Kalman Filter to forecast the next “m” measurements. During the Kalman-Filtering interval, error corresponds to the difference between a-priori predicted measurements and true measurements at each time-step. Likewise, error during the final stage corresponds to the difference between true and forecasted measurements. We have previously derived and simplified the full analytical gradients of this process ([27]) which are fed into a stochastic gradient/Hessian optimization algorithm to update parameter estimates. In the present paper, we used Nesterov-Accelerated Adaptive Moment Estimation (NADAM; [60]) for the gradient updates, however a wide-variety of gradient-based algorithms are included with the publicly available code.

For brevity, we present the gBPKF equations ([27]) in their most general form with all parameters contained within the (high-dimensional) variable *θ*. We also focus upon regressing gradients through the Kalman-Filtering stage, as opposed to the full filtering-forecasting sequence as the forecasting stage is simply a special case of the Kalman Filter with the probabilistic (*P*) and update (*K*) terms removed. Our objective is to minimize a function ℒof prediction error 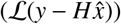. In the current paper, we sued the smoothed Huber-loss 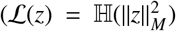, see Sec. 8.3), For a set of initialization times *t*_0_ and filter-length *m*, we denote the accumulation of error from step *t* up to *m* as:

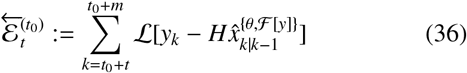

The full objective is to solve for *θ* which minimizes the total error over all initialization times 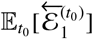. For brevity’s sake, we omit the notation indicating dependencies upon parameters (*θ*), previous measurements (ℱ [*y*]) and start-times. Unlike a conventional dynamical systems model, the Kalman Filter evolves both the state-expectation 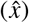 and error-covariance *P*. To eliminate redundant equations, we use *ω* to denote the combined set 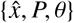 with time indices on 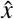, *P* being all previous time-steps.

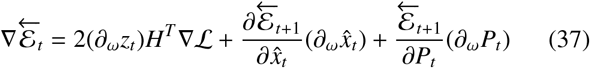

Evaluating the first term: 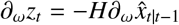 and backpropagating through the Kalman update:

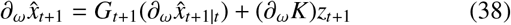

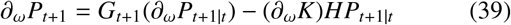

Fixing *H, Q, R*, the Kalman gain is a direct function of *P*_*t*|*t*−1_ (and its influence on S). Using that *K*_*t*_ minimizes *Tr*[*P*_*t*_] and applying the implicit function theorem:

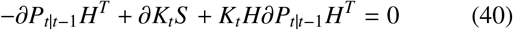

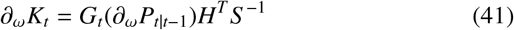

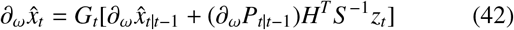

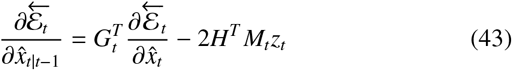

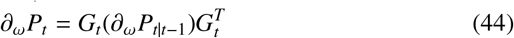

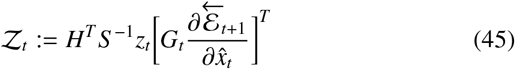

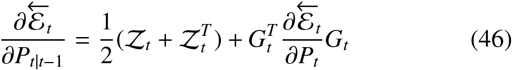

We are now ready to propagate errors through the a-priori statistics which, depending upon the choice of Filter, can be estimated in various ways. The following equations hold equivalently for the exact statistics and any reasonable means of approximating them as implemented in any currently used Kalman Filter. Specifically, we only assume that the approximations of 𝔼 preserve linearity and *cov* preserve bilinearity. We therefore have the general recursions, for 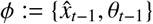:

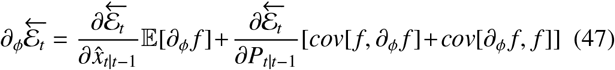

The gradients with respect to *P*_*t*_, however, are specific to the filter choice, although general forms are presented in [27]. For the EKF ([28]), as used in the current work, we have:

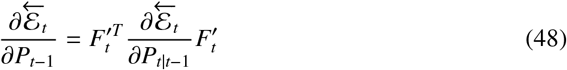

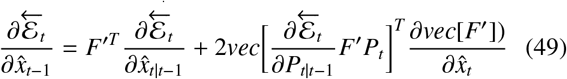

The final gradient of total error with respect to parameter for the filtering stage is:

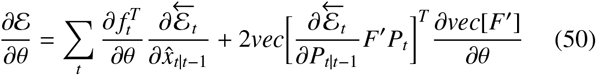

Gradients with respect to sample-based filters (e.g., Unscented Kalman Filter, [29]) can be found in [27]. In the generative and forecasting stages, gradients follow the normal back-propagation-through-time recursions as there is no Kalman-filtering.

### 8.5. Estimating Nonlinear Posteriors

In general, our approach (gBPKF) is compatible with a wide variety of techniques for estimating posterior means and covariances following a nonlinearity (see [27]). In the current work, however, we use a simple batch formulation of the Extended Kalman Filter (EKF; [28]) which allows the Kalman Filter to run on many data segments in parallel. We use 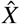 to denote the concatenated state estimates for all initial time-points within a minibatch, with a common error-covariance 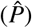 used for all members of the minibatch. We use the set-valued index index {**q**} to denote the current time step for all data segments within the minibatch. The prior distribution and posterior statistics are:

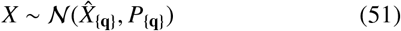

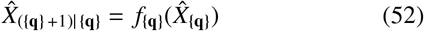

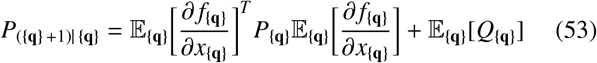

Hence we alter the EKF to use the average Jacobian over the minibatch. This step allows many data segments to be run in parallel by sharing a common covariance *P*_{**q**}_ at the cost of having less sensitivity to time-variation in the nonlinearity. To counteract this drawback, we chose minibatches as temporal chunks (80 initial timepoints with 5-step=10ms spacing) so as to track nonstationary statistics. In all of our applications, the process noise *Q*_{**q**}_ did not depend upon time, so the rightmost expectation simplifies to *Q*.

### 8.6. Generating the Connectivity Mask

We constrained eligible inter-area connections using a liberally-defined connectivity mask which was generated based upon previous modeling with fMRI data ([19]) as described below. While this step is not an inherent requirement of our approach, it is important for ensuring that fits are robust with M/EEG data. Specifically, this constraint can counteract modest deviations in the forward model (i.e., head position relative sensors) and promotes well-posedness when multiple brain areas have weak contributions to the M/EEG signal. We formed our connectivity-mask based upon group-level consensus from the fMRI MINDy models ([19]) using the union of several criteria to avoid reliance upon any single measure of what constitutes a non-trivial connection. This procedure retains sensitivity to small, but consistent connections as well as connections that may not show up in every subject but are (on average) large. We rescaled each fMRI connection matrix (two runs per subject) to have a root-mean-square value of 1, excluding self-connections. For thresholding, we used an absolute magnitude threshold of applied to each matrix. We also note that, unlike the M/EEG models presented in the current work, the fMRI MINDy models use a single connection per region-pair which can be positive or negative as opposed to both EE and EI projections. We then generated the connectivity mask through the union of three criteria:

1. Average magnitude: We admitted the top 15% of connections in terms of magnitude for the group-average.
2. Consensus: We admitted connections with an average post-threshold sign greater than 0.8 or less than -0.6 across subjects/runs.
3. Minimal Count: For each parcel we admitted the (up-to) three largest positive and negative connections in terms of input, output, and symmetrized-strength (average of input and output) with overlap between these categories.

We then symmetrized the resultant matrix so that if *W*_*i*, *j*_ is an admissible connection, so is *W*_*j,i*_. The final mask had 2,522 admissible long-distance connections out of the possible 9,900 (25.2%). We used the same connectivity mask to constrain long-distance excitatory-excitatory connections and long-distance excitatory-inhibitory connections for a total of 5,044 inter-region conncetions.

### 8.7. Promoting Unique Solutions

A key trap in data-driven modeling with latent-variables is that several model solutions may be observationally-equivalent, meaning that they behave identically in terms of the measured variables even if they are nonidentical in terms of the non-measured variables. In our case, this ambiguity largely corresponds to models in which the predicted excitatory activity is the same, but not the inhibitory activity which lacks a predefined unit (scale) since it is not directly measured. For many use-cases this ambiguity is irrelevant as it only affects interpretation of the absolute scale for inhibitory activity/parameters. However, for good form, we reduce the parameter-space to ensure unique solutions. To be clear, the model is not overparameterized in the usual sense–for a fixed set of latent states {*x*} the optimal parameter choices are well-posed. Rather, the difficulty arrives with the possibility of transformed systems behaving identically in terms of measurements. This relationship is called observational equivalence and, in the deterministic case, can be stated as follows: the systems *x*_*t*+1_ = *f*_*t*_(*x*_*t*_) and 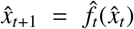 are observationally-equivalent with respect to the measurement-process *y*(*x*) = *Hx* if for every *x*_0_ there exists 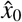 s.t. 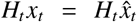 with *x* and 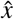 evolving according to *f* and 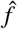, respectively. As a specific case, consider *f* (*x*) = *Wψ*(*S x* + *v*) + *Dx* + *c* with *S* and *D* diagonal. If the vectors *b, q* satisfy *H*_*t*_*diag*(*b*) = *H*_*t*_ and *H*_*t*_*q* = 0, ∀*t*, then the transformed system 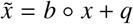 is observationally-equivalent to *x* and has parameters:

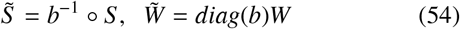

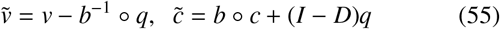

Here and in all later cases we use to denote element-wise multiplication (Hadamard product). The inverse-notation *b*^−1^ is understood to be applied element-wise for vectors. In the above case, we note that *b, q* may be chosen arbitrarily for inhibitory indices since the corresponding measurement gains are zero (*j* inhibitory implies *H*_*i*, *j*_ = 0). Thus there are at least two arbitrary degrees of freedom per-region in the unrestricted model: parameter choices can be altered such that they uniformally shift or rescale estimates of the inhibitory population activity. To remove these arbitrary degrees of freedom, we fixed parameters of the inhibitory nonlinearity to be *S* _*I*_ = 1, *V*_*I*_ = 0, thereby removing the shift and scale symmetries, respectively.

However, some invariances may remain due to volume conduction and referencing. These invariances do not affect temporal dynamics in the measurement space, and instead reflect translating the system in a direction to which the M/EEG sensors are blind (i.e., a pattern of brain activity in which electric fields cancel at the sensor-level). In mean-referenced EEG, for instance, the relativistic nature of voltages mean that the result of shifting each region’s activation by a constant (i.e. 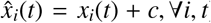) will be observationally-equivalent to the original system.

To further constrain the problem, we reduced the space of recurrent connectivity patterns by assuming that they shared a common non-negative spatial gradient (*b*) within-subject and that each recurrent connectivity vector 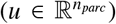 can be expressed as an affine transformation of this gradient:

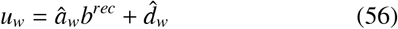

with separate scalar values of *a*_*w*_, *d*_*w*_ for *w* ∈ {*EE, EI, IE, II*}. In experimentation, we found that this restriction can be further eased with separate excitatory and inhibitory spatial gradients, but do not recommend fully unrestricting the recurrent connections due to the aforementioned possibility of parameter symmetries. The associated software enables users to define arbitrary constraints of this form between parameters and use multiple-component bases (matrix-valued *b*^*rec*^, vector-valued *a*_*w*_).

### 8.8. Defining the MEG Measurement Model

For this initial validation with MEG, we built a measurement model in which the timeseries had already been source-localized with Minimum Norm Estimation (MNE), while the optimization objective was calculated in a rank-reduced subspace of this projection (described below). The noise-covariance matrix was adapted from empty-room recordings (*R*^*chan*^). Separately for each scan, we rank-reduced the data according to the singular-values of the post-ICA leadfield: 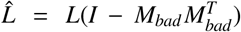. We rank reduced *L* based upon a singular-value threshold of 1% the maximal singular value and denote the reduced left-singular vector matrix *U*_*L*_. We projected the leadfield onto this space (premultiplied by 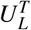) which, inherently removes the ICA-censored dimensions from the leadfield (reflecting the fact that those dimensions were discarded). Source-estimation was then performed using MNE on this new leadfield to produce the source-inversion matrix ℳ_0_. We rescaled rows of the MNE inverse solution (ℳ_0_) so that 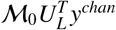 had unit variance for each channel. Thus, the final inverse transformation was:

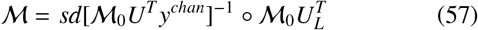

We correspondingly transformed variables such that the Kalman-Filter receives source-estimates as the measurement variable:

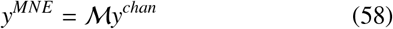

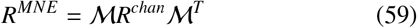

Within this space, the measurement matrix was:

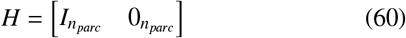

reflecting a direct, but noisy, source-estimate for excitatory activity and no sensitivity to inhibitory activity. Thus, standard source-estimates were fed into the Kalman Filter during state-estimation. However, the model error was assessed over the more parsimonious space span(ℳ) which removes artificial dimensions created during source-estimation. Thus, models are not held to fully obey the source estimates in dimensions of ambiguity. Writing the singular-value decomposition of ℳ:

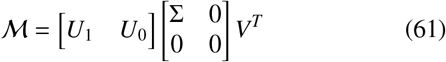

We used the cost matrix: 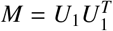 (see Eq. 28).

### 8.9. gBPKF Training Parameters for MEG Data

The gBPKF algorithm ultimately consists of training a (biological) recurrent neural network with the Kalman-Filter as an additional, interconnected unit. Thus, the hyperparameter classes are conceptually analogous to any deep learning scenario although the actual training equations are quite different. For the HCP MEG data, we used a filtering period of 26-steps with the Kalman correction only applied every fifth step starting at one (i.e., 6 updates total) followed by a forecasting stage of 35 prediction steps. On each batch, initial distributions (mean and covariance) were estimated by simultaneously simulating 50 initial conditions with process noise for 50 time-steps. Initial values of *x* for these simulations were drawn from a moving record of predictions during previous forecasting stages. This step reduces the time to reach steadystate (*t* → ∞) distributions, thereby enabling shorter simulation periods. The covariance of this simulation was smoothed according to an autoregressive average with coefficient 0.05, i.e. *cov*(*k*) = .95*cov*(*k* − 1) + .05*cov*[*sim*(*k*)]. Gradients were fully propagated through each simulation’s influence on *cov*[*sim*(*k*)], but not to previous training iterations (which had different parameter estimates). Minibatches consisted of 80 time segments and there were 4 minibatches per training iteration. To improve memory-management we sampled initial time-points in chunks, with 5-step = 10ms spacing between initial time-points, so that the measurement timeseries overlapped for neighboring initial time-points within a minibatch. This choice also retains sensitivity to non-stationary statistics that would otherwise be lost in batch-EKF (Sec. 8.5). For efficiency, minibatches began at the filtering stage, so state distributions were estimated once per training iteration instead of once per minibatch. Training used a fixed budget of 150,000 iterations (determined based upon examination of convergence rate). Gradients were clipped ([61]) with a moving threshold of 3 times the average norm over the past 200 iterations. Parameter updates were performed using the NADAM algorithm ([60]) with hyperparameters *β*_1_ = .98, *β*_2_ = .99, *ν* = .0001 and rate .0001.

### 8.10. Ground-Truth Simulations

In previous work, we validated and benchmarked the gBPKF algorithm in estimating parameters for unstructured, randomly generated recurrent neural networks ([27]). However, these previous analyses did not consider whether our approach would perform equally well with networks obeying an excitatory-inhibitory structure. We therefore performed a new set of ground-truth simulations which directly mirrored our model setup. Each ground-truth model consisted of interconnected “regions” which each contained an excitatory and an inhibitory population. Only excitatory populations made long-distance connections which targeted both excitatory and inhibitory populations. Measurements were simulated by multiplying excitatory activity with a randomly generated “leadfield” and adding temporally-independent Gaussian noise (with covariance *R*).

To parameterize the ground-truth models, we first randomly generated a symmetric connectivity mask *W*_*mask*_ with 25% density on the off-diagonals and zero on the diagonals. The same connection mask is used for long-distance EE and EI connections. The EE and EI connection strengths are also generated from similarly constructed distributions, hence we omit the EE vs. EI distinction and refer to a single connectivity matrix for now. Other than symmetry 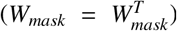, the entries of *W*_*mask*_ are uncorrelated. As with data applications, the values of *W*_*mask*_ simply indicate if a connection is plausible while the actual value could be effectively zero. Connection strengths were not symmetric and were generated as the sum of a sparse matrix and a low-rank matrix. The sparse matrix *W*_*s*_ was distributed:

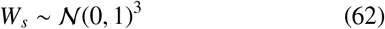

The low rank component (*W*_*L*_) was the product of two rectangular matrices 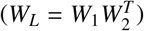 generated according to:

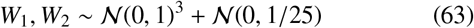

These terms are combined according to

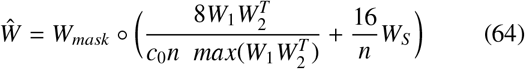

Note that the individual connection strengths are inversely proportional to the number of nodes (*n*) so that the total input strength is similar across simulation sizes (akin to subdividing the brain into smaller pieces). This process was used to generate long-distance EE connections 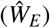 and EI connections 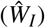. For the EE connections *c*_0_ = 2, whereas for the EI connections *c*_0_ = 1.5, hence EE projections were generally stronger than EI. Recurrent connections were generated from a narrower distribution designed to sustain nontrivial dynamics (oscillations etc.). The base values for each local connections were: EE: .5, EI: 1.25, IE: -1.25, II: -.25. The spatial gradation in local connectivity (*b*) was generated as *b* ∼.85 + .3𝒩(0, *I*_*n*×*n*_)^2^. The final local connectivity strength was generated by multiplying the base value for each connection type by *b*. As simulations were randomly-parameterized, a small number produced pathological steady-state behavior in which model inversion was theoretically impossible. Specifically, in these cases the dynamics generated attractors so far outside the dynamic range of *ψ* that the network was stuck in a saturated state. These dynamics are unambiguous and easy to detect. We removed simulations in which the median moving-variance of *ψ* was less than .01 for at least half of the populations/nodes (normal values are around .9 for all nodes).

